# Plastome phylogenomic study of Gentianeae (Gentianaceae): widespread gene tree discordance and its association with evolutionary rate heterogeneity of plastid genes

**DOI:** 10.1101/2020.04.02.021840

**Authors:** Xu Zhang, Yanxia Sun, Jacob B. Landis, Zhenyu Lv, Jun Shen, Huajie Zhang, Nan Lin, Lijuan Li, Jiao Sun, Tao Deng, Hang Sun, Hengchang Wang

## Abstract

**Background:** Plastome-scale data have been prevalent in reconstructing the plant Tree of Life. However, phylogenomic studies currently based on plastomes rely primarily on maximum likelihood (ML) inference of concatenated alignments of plastid genes, and thus phylogenetic discordance produced by individual plastid genes has generally been ignored. Moreover, structural and functional characteristics of plastomes indicate that plastid genes may not evolve as a single locus and are experiencing different evolutionary forces, yet the genetic characteristics of plastid genes within a lineage remain poorly studied.

**Results:** We sequenced and annotated ten plastome sequences of Gentianeae. Phylogenomic analyses yielded robust relationships among genera within Gentianeae. We detected great variation of gene tree topologies and revealed more than half of the genes, including one (*atpB*) of the three widely used plastid markers (*rbcL, atpB* and *matK*) in phylogenetic inference of Gentianeae, are likely contributing to phylogenetic ambiguity of Gentianeae. Estimation of nucleotide substitution rates showed extensive rate heterogeneity among different plastid genes and among different functional groups of genes. Comparative analysis suggested that the ribosomal protein (RPL and RPS) genes and the RNA polymerase (RPO) genes have higher substitution rates and genetic variations in Gentianeae. Our study revealed that just one (*matK*) of the three (*matK, ndhB* and *rbcL*) widely used markers show high phylogenetic informativeness (PI) value. Due to the high PI and lowest gene-tree discordance, *rpoC2* is advocated as a promising plastid DNA barcode for taxonomic studies of Gentianeae. Furthermore, our analyses revealed a positive correlation of evolutionary rates with genetic variation of plastid genes, but a negative correlation with gene-tree discordance under purifying selection.

**Conclusions:** Overall, our results demonstrate the heterogeneity of nucleotide substitution rates and genetic characteristics among plastid genes providing new insights into plastome evolution, while highlighting the necessity of considering gene-tree discordance into phylogenomic studies based on plastome-scale data.

## Background

Whole plastomes have become more accessible with the explosive development of next-generation sequencing (NGS) technologies [1-3]. Due to the unique mode of inheritance, conservativeness in gene content and order, and high copy number per cell [4, 5], plastomes have been widely used in reconstructing the plant Tree of Life (e.g. [6-10]). Moreover, compared to standard fragment DNA barcodes, plastome-scale data can provide an abundance of informative sites for phylogenetic analyses [5, 11-13]. Nonetheless, the effectiveness of plastome-scale data is ultimately reflected by the extent to which they reveal the “true” phylogenetic relationships of a given lineage [14].

Although plastomes have been canonically regarded as a linked single locus due to its uniparental inheritance and lack of sexual recombination [4, 5, 15], structural and functional characteristics of plastomes suggest that plastid genes may not evolve as a single locus and are experiencing different evolutionary forces [16-18]. Furthermore, despite evolving at a lower rate than genes in the nucleus [19], rate variation within the plastomes has been documented between the single copy and inverted repeat regions, different functional groups of genes, or across angiosperm lineages (e.g. [16, 20-22]). Factors contributing to rate variation and affecting the evolution of different plastid genes include mutation rate variation across families and between coding/non-coding regions, as well as variation in the SSC due to the presence of two configurations of the inversion [14]. However, in many recent phylogenomic studies using the full plastomes, only the results from the full concatenated data set are presented (e.g. [6, 7, 23]). In these cases, gene-tree discordance due to evolutionary rate variation of individual genes remains poorly understood. In addition to concatenated approaches, multispecies coalescent methods (MSC) account for gene tree heterogeneity allowing for the assessment of ancient hybridization, introgression, and incomplete lineage sorting (ILS) by using the summed fits of gene trees to estimate the species tree [24, 25]. Recently, phylogenomic studies suggest that a combination of concatenated and coalescent methods can produce accurate phylogenies and benefit the investigation into the incongruence between gene trees and species trees [18, 26].

Comparative genomic studies based on plastomes have mainly concentrated on structure variations, such as contraction or expansion of inverted repeats (IR) (e.g. [27-29]) and genomic rearrangements (e.g. [30-33]), yet the genetic characteristics of plastid genes within a lineage, such as genetic variation and phylogenetic informativeness, remain poorly studied. These characteristics may vary among different genes or functional groups of genes and are of great importance in our understanding of plastome evolution and phylogenetic inference. Additionally, the correlation between evolutionary rate and gene characteristics can be invoked as an explanation of the primary impetus of plastome evolution [16, 20, 34, 35].

The tribe Gentianeae, with its two subtribes Gentianinae and Swertiinae, include ca. 950 species in 21 genera, exhibiting the highest species diversity of the Gentianaceae [36]. Members of Gentianinae are easily distinguishable from Swertiinae by the presence of intracalycine membranes between the corolla lobes and plicae between the corolla lobes, with both traits absent in Swertiinae [36-38]. Although several phylogenetic studies have confirmed the monophyly of both subtribes [36-40], the generic delimitation within Gentianeae remains ambiguous, especially within Swertiinae, with some genera (e.g., *Swertia* L., *Gentianella* Moench, *Comastoma* (Wettst.) Toyok., *Lomatogonium* A.Braun) being paraphyletic [38]. The current phylogeny of Gentianeae is based upon a few DNA markers, commonly including ITS, *atpB, rbcL, matK*, and *trnL–trnF* [36-41], thus a full taxonomic and evolutionary understanding of these groups is hindered by the unsatisfactory phylogenetic resolution.

To gain new insights into the evolution of plastomes, and to improve delineation of the phylogenetic affinities among genera in Gentianeae, we constructed a dataset of plastome sequences including 29 Gentianeae species and three outgroups. We generated 76 protein-coding gene (PCG) sequences to infer phylogenies via both concatenated and coalescent methods, while also characterizing genetic features of plastid genes. Our specific goals are to (a) test whether plastome-scale data is effective in resolving enigmatic relationships within Gentianeae; (b) investigate characteristic diversity of plastid genes of Gentianeae; and (c) explore the correlation of evolutionary rate heterogeneity with gene characteristics as well as gene-tree discordance.

## Results

### Characteristics of Gentianeae plastomes

A total of ten species were newly sequenced (Supplementary Table S1) representing ten genera (seven newly reported) of Gentianeae. After *de novo* assembly, we generated a single contig for each newly sequenced plastome. The mean sequencing coverage ranged from 646× (*Gentiana urnula*) to 3,538× (*Gentianopsis paludosa*). All ten plastomes display the typical quadripartite structure composed of a large single copy (LSC), a small single copy (SSC), and two inverted repeats (IRa and IRb). The length of the ten plastomes range from 139,976 bp in *Kuepferia otophora* to 153,305 bp in *Halenia elliptica* (Table 1). All the plastomes have four rRNAs and 30 tRNAs and are in the same gene order (Figure 1). Gene loss involving *ndh* genes in the genus *Gentiana* was detected. Moreover, the *rpl33* gene was found to be lost in *Comastoma pulmonarium* and *Swertia hispidicalyx* (Figures 1, 2 and S1). Plastomes of Gentianeae were highly conserved with only one event of IR expansion occurring in *Halenia elliptica*, where IR regions expanded to the *rpl22* gene.

**Table 1.**
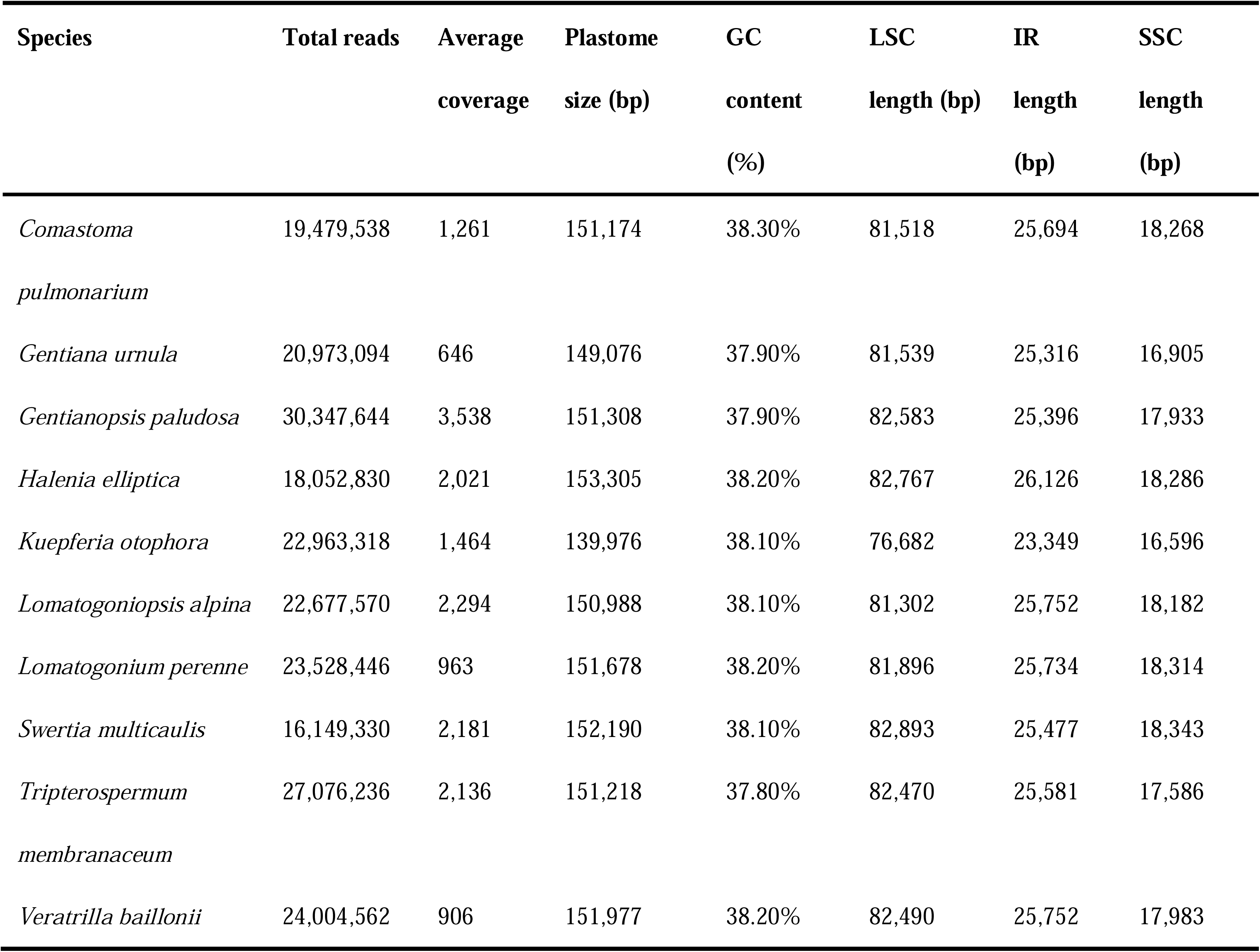
Plastome features of newly sequence Gentianeae species. Abbreviations: LSC, large single copy; SSC, small single copy; IR, inverted repeat.

**Figure 1.**
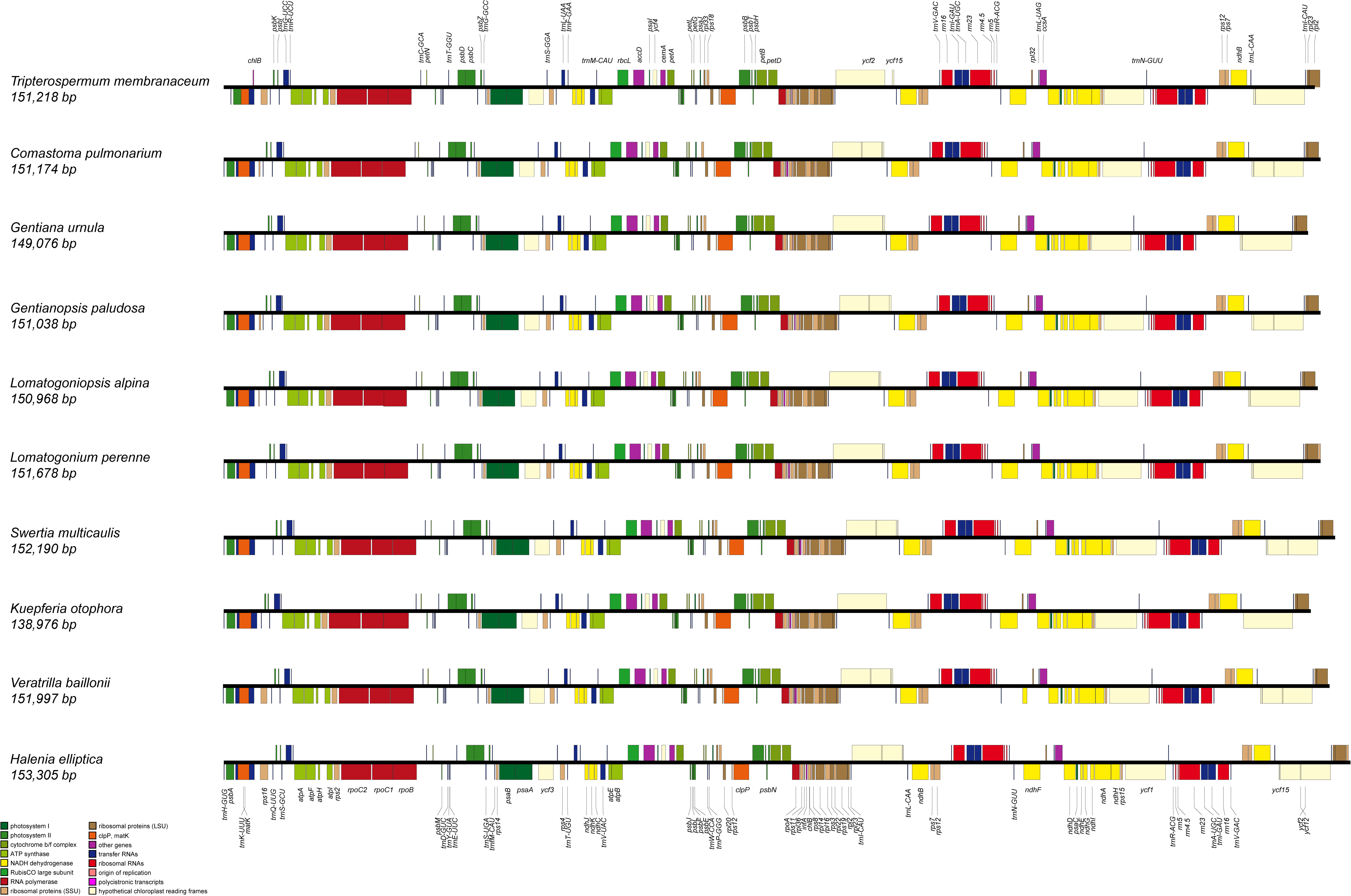
Linear maps of the newly sequenced plastomes of Gentianeae. Full-color boxes with labeled gene names highlight coding sequences by gene class, as summarized to the bottom-left corner.

**Figure 2.**
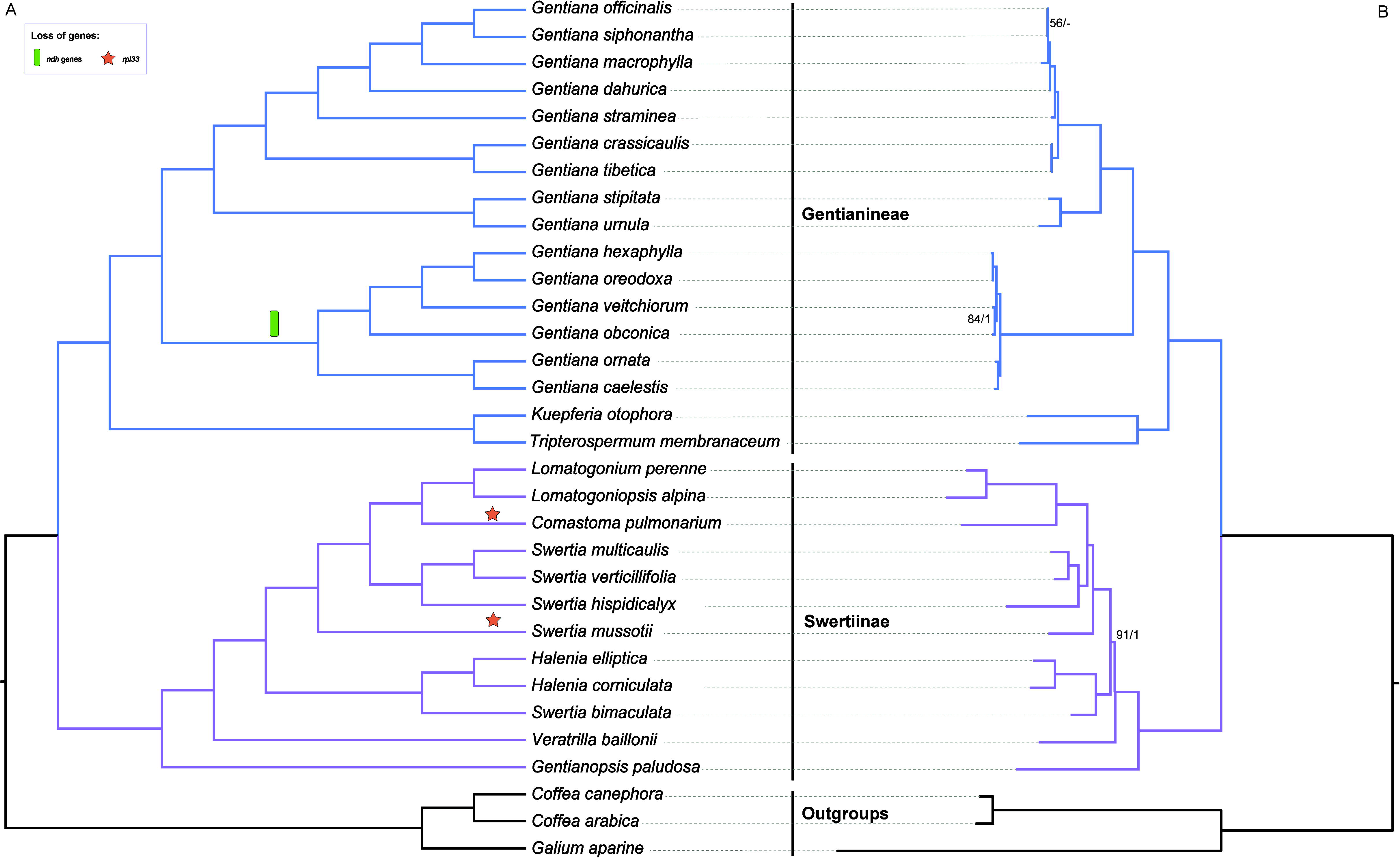
Phylogenetic result of Gentianeae, with **(A)** distribution of gene loss mapped on the cladogram generated by coalescent method in ASTRAL-III. Local posterior probabilities (PP) of all branches were 1.0, and were not showed in the tree. **(B)** Maximum likelihood phylogram of Gentianeae from partitioned concatenated matrix of 76 plastid protein-encoding genes using RAxML. Maximum likelihood bootstrap (BS) values and the PP calculated from MrBayes are shown at nodes, except nodes with 100% BS and 1.0 PP, ‘-’ indicates no support value.

### Phylogenetic relationships within Gentianeae

The concatenated alignment of the 76-gene, 32-taxa dataset had 69,579 bp in length consisting of 9,228 parsimony-informative sites. Our phylogenomic analyses improved the resolution and robustness of affinities among genera in Gentianeae, with most clades exhibiting high support values. For concatenated data set, partitioned Maximum likelihood (ML) and Bayesian Inference (BI) analyses (Figure 2) yielded identical tree topologies with unpartitioned data sets (Figure S2). The same tree topology was also inferred with the coalescent analysis (Figure 2). The monophyly of two subtribes, i.e. Gentianinae and Swertiinae, were supported in all analyses. In Gentianinae, a clade consisting of *Tripterospermum* and *Kuepferia* was sister to *Gentiana*. Within *Gentiana*, species exhibiting *ndh* gene loss events formed a distinct clade and were sister to other members of *Gentiana*. In Swertiinae, *Swertia* was revealed as nonmonophyletic due to the close relationship between *Swertia bimaculate* with the monophyletic genus *Halenia*.

### Gene trees landscape

We employed Principal Coordinate Analysis (PCoA) of the unrooted gene trees inferred with ML and the species trees estimated from the concatenated and coalescent analyses to investigate incongruence between species trees and gene trees [42]. In our analysis, the first and second axes of the PCoA explained 13.8% and 4.7% of the variation in tree topologies, respectively. The species trees inferred with the two different methods clustered together, while gene trees showed greater variation. We calculated the distance between gene trees and the coalescent species tree to represent gene-tree discordance (GD) of each gene (Mean: 14.789; median: 14.560). Among 75 genes tested, *rpoC2* had the lowest GD value (GD = 1.54E-05) and *petL* had the highest (GD = 35.626). Gene trees from the three traditionally used plastid genes (*rbcL, atpB* and *matK*; GD = 2.213, 2.029 and 0.928) were close to the species trees (Figure 3A, Table S2).

**Figure 3.**
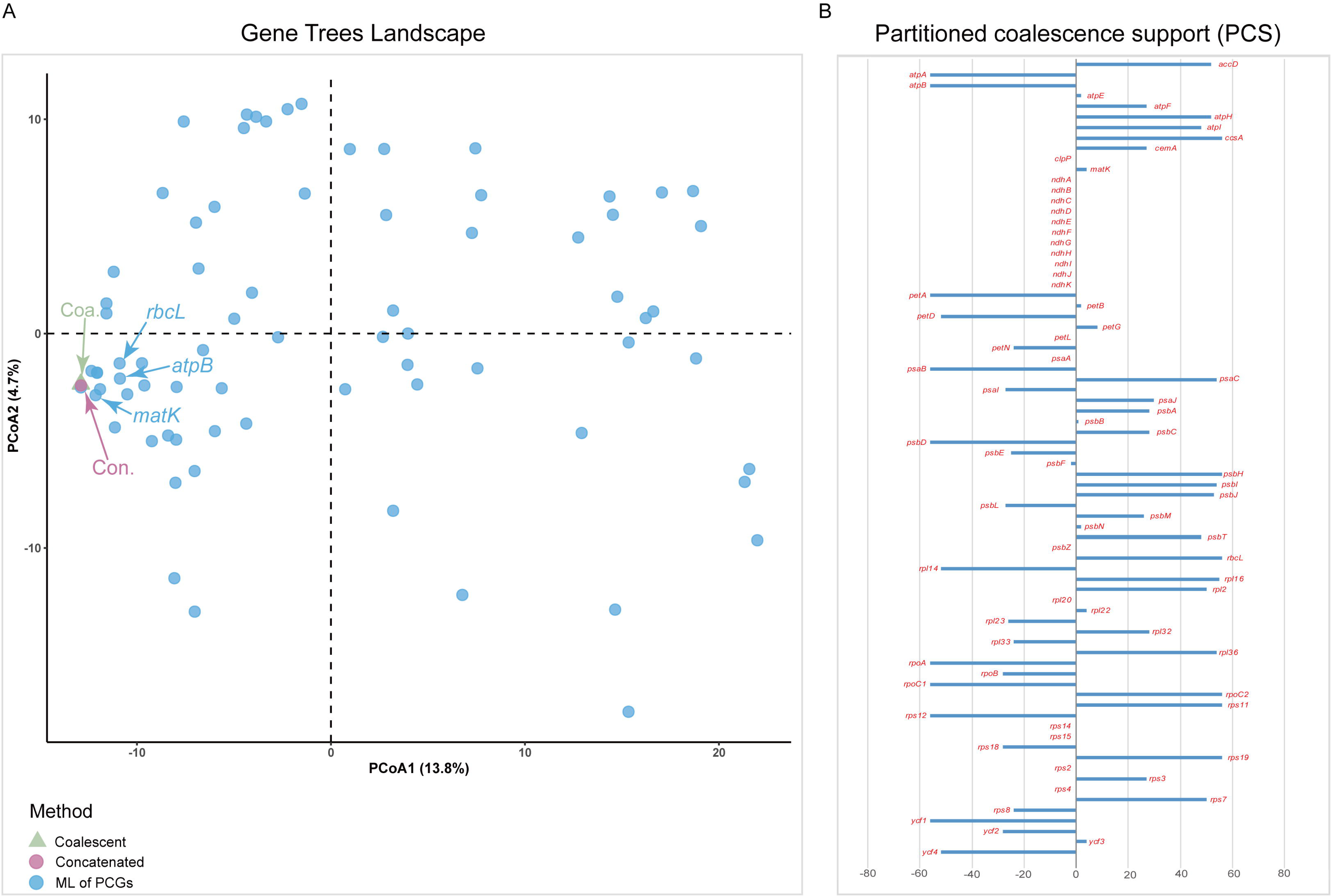
Discordance of plastid gene trees. **(A)** Principal coordinate analysis depicting ordinations of two species trees (green and pink) versus 75 plastid protein-coding gene (PCG) trees (blue) using unrooted Robinson-FouldS algorithms. Since TREESPACE only accepts groups of trees containing the same tips, *rpl33* locus was removed. Genes widely used in phylogenetic studies of Gentianeae are indicated in the plots (matK, rbcL, and atpB). **(B)** Partitioned coalescence support (PCS) of each PCG. Names of 76 PCGs are labeled in red color.

We also computed the partitioned coalescence supports (PCSs) of the 76 PCGs. PCS can be positive, negative, or zero, indicating support, conflict, or ambiguity, respectively [43]. The results revealed 33 PCGs with positive PCS scores, 23 PCGs with negative PCS scores and 20 PCGs with zero PCS scores (Figure 3B, Table S2). Six PCGs (*ccsA, psbH, rbcL, rpoC2, rps11* and *rps19*) were estimated with highest PCS score (PCS□=□56). Among the three widely used plastid markers in previous phylogenetic studies of Gentianeae, *rbcL* and *matK* had positive PCS scores, whereas *atpB* had a negative PCS score.

### Nucleotide substitution rates

Synonymous (d*S*) and nonsynonymous (d*N*) substitution rates, along with d*N*/d*S* were estimated for the 76 PCGs to detect evolutionary rate heterogeneity and to represent different selection regimes acting on PCGs (Table S2). Among the 76 genes, *rps22, rps15* and *rpl32* had relatively higher d*S* values, and *ycf1, matK* and *rpl33* had higher d*N* values (Figure 4A, Table S2). All 76 PCGs exhibited considerably low values of d*N*/d*S*, indicating that they have been under purifying selection. We also compared evolutionary rates among nine functional groups and one group of other genes (OG, Table 2). The OG had the highest median values of d*N* and d*N*/d*S* but a moderate d*S* median value. Genes encoding subunits that are integral to photosynthetic processes, such as ATP synthase (ATP), cytochrome b6f complex (PET), and photosystems I and II (PSA and PSB), generally have lower rates of nucleotide substitution than other functional groups of genes. The RNA polymerase (RPO) genes showed highly accelerated rates in d*N* and d*N*/d*S*, while genes encoding proteins of the ribosomal large subunit (RPL) had the highest d*S* value (Figure 4B). We also concatenated the genes located in the LSC, SSC and IR to investigate substitution rate differences among IR vs SC regions. The IR region had the lowest d*N* and d*S* values (0.095 and 0.200 respectively), while the SSC region had the highest (d*N*= 0.464; d*S*= 0.958), followed by the LSC region (d*N*= 0.134; d*S*= 0.878).

**Table 2.** Plastid genes and functional groups included in analyses.

| Functional groups | Genes |
| --- | --- |
| Photosystem I (PSA) | *psaA, psaB, psaC, psaI, psaJ* |
| Photosystem II (PSB) | *psbA, psbB, psbC, psbD, psbE, psbF, psbH, psbI, psbJ, psbL, psbM, psbN, psbT, psbZ* |
| Cytochrome B<sub>6</sub>f complex (PET) | *petA, petB, petD, petG, petF, petN* |
| ATP synthase (ATP) | *atpA, atpB, atpE, atpF, atpH, atpI* |
| Rubisco large subunit (Rubisco) | *rbcL* |
| RNA polymerase (RPO) | *rpoA, rpoB, rpoC1, rpoC2* |
| Ribosomal proteins large subunit (RPL) | *rpl2, rpl14, rpl16, rpl20, rpl22, rpl23, rpl32, rpl33, rpl36* |
| Ribosomal proteins small subunit (RPS) | *rps2, rps3, rps4, rps7, rps8, rps11, rps12, rps14, rps15, rps18, rps19* |
| NADH dehydrogenase (NDH) | *ndhA, ndhB, ndhC, ndhD, ndhE, ndhF, ndhG, ndhH, ndhI, ndhJ, ndhK* |
| **Other genes (OG)** |  |
| Conserved coding frame | *ycf1, ycf2, ycf3, ycf4* |
| Acetyl-CoA-carboxylase | *accD* |
| ATP-dependent protease | *clpP* |
| Cytochrome c biogenesis | *ccsA* |
| Membrane protein | *cemA* |
| Maturase | *matK* |

**Figure 4.**
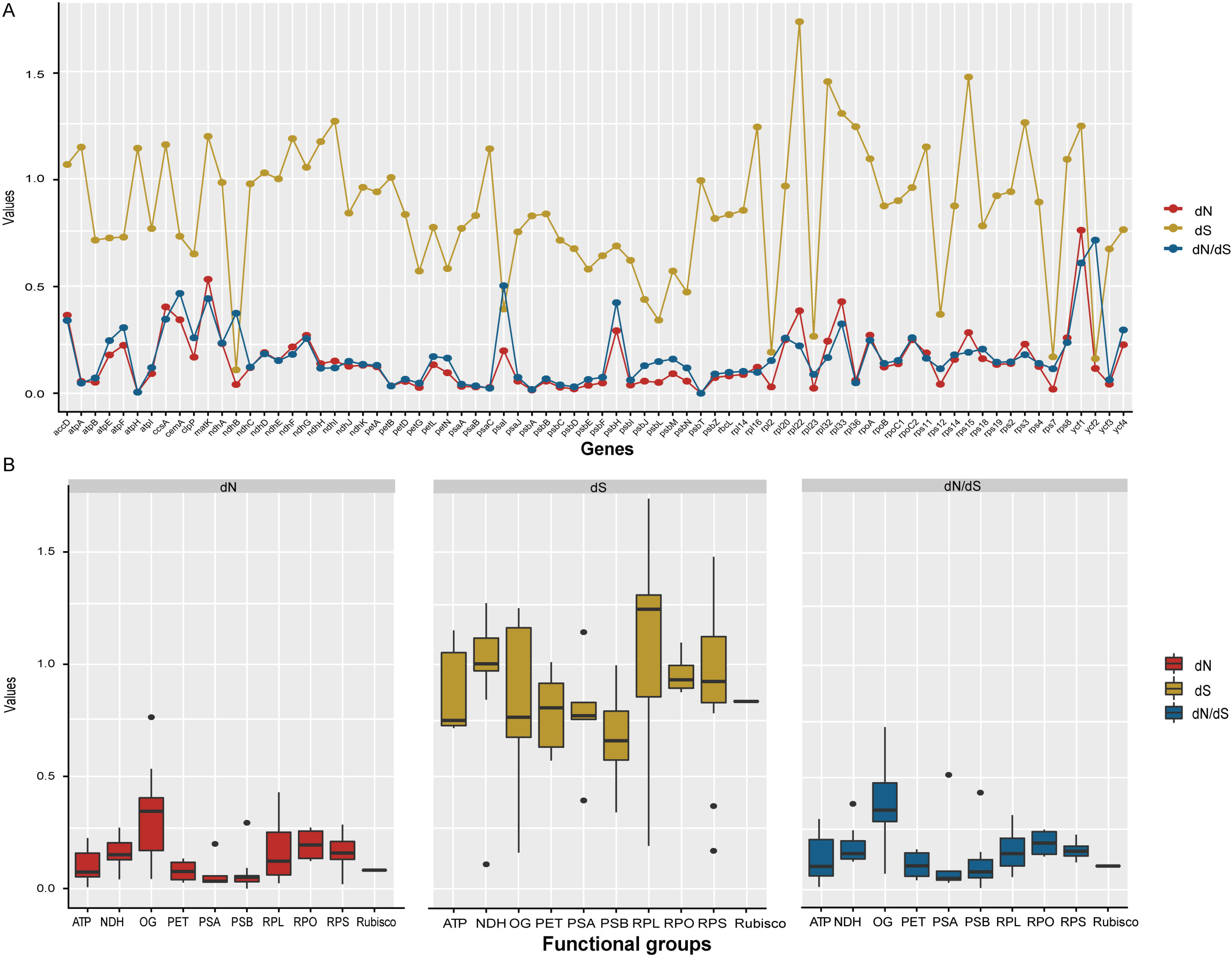
The estimations of nonsynonymous (d*N*), synonymous (d*S*) substitution rates and d*N*/d*S* of **(A)** plastid protein-coding genes (PCG), and of **(B)** functional groups of genes. Detailed information of functional group is provided in **Table 2**.

### Genetic characteristics of plastid genes

We calculated the nucleotide diversity (π) and percent variability (PV) to represent genetic variation of PCGs (Table S2). The values of π ranged from 0.0077 (*rps7*) to 0.0884 (*ycf1*), and values of PV ranged from 0.0271 (*ndhF*) to 0.3872 (*rpoC1*, Figure 5A, Table S2). Among the functional groups, RPO, RPS, RPL and NDH showed both high nucleotide diversity and percent variability (Figures 5B and C), especially genes in the RPO group. Notably, RPO had highest median value of gene length (Figure 5D). The net phylogenetic informativeness (PI) for the 76 PCGs used in phylogenetic analysis were measured using PhyDesign (Figure S2; Table S2). The *ycf1* gene had the highest net PI of all PCGs, followed by *rpoC2, ndhF* and *matK*. Genes with high PI were also genes with longer length, suggesting a large contribution of gene length to phylogenetic informativeness. Relatively conservative genes with less phylogenetic informativeness were primarily associated with photosynthesis and were shorter in length (< 200 bp).

**Figure 5.**
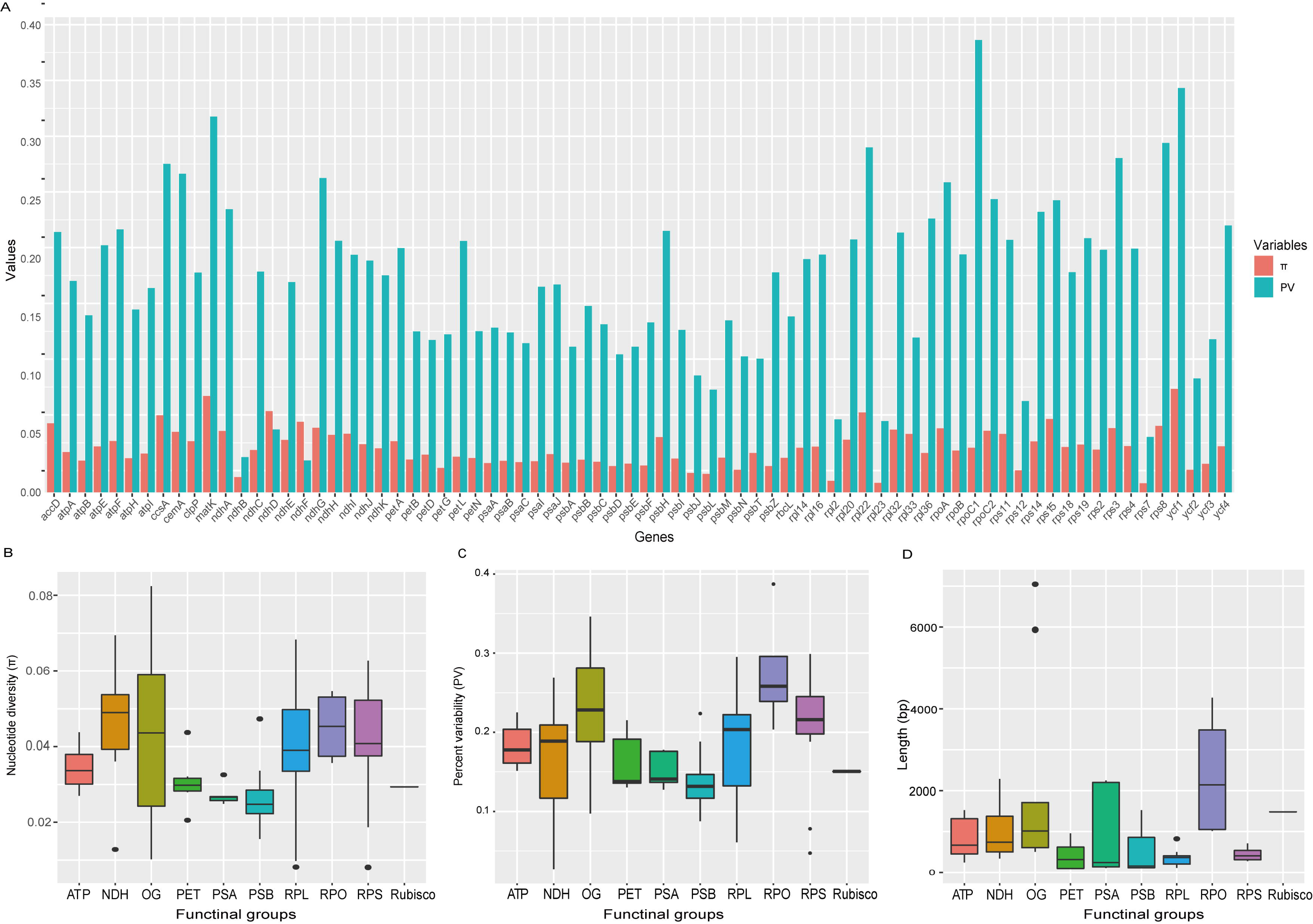
Genetic variation among plastid protein-coding genes (PCGs) and among functional groups. **(A)** Nucleotide diversity (π) and percent variability (PV) for 76 PCGs. **(B)** Nucleotide diversity (π), **(C)** percent variability (PV) and **(D)** gene length for functional groups.

**Figure 6.**
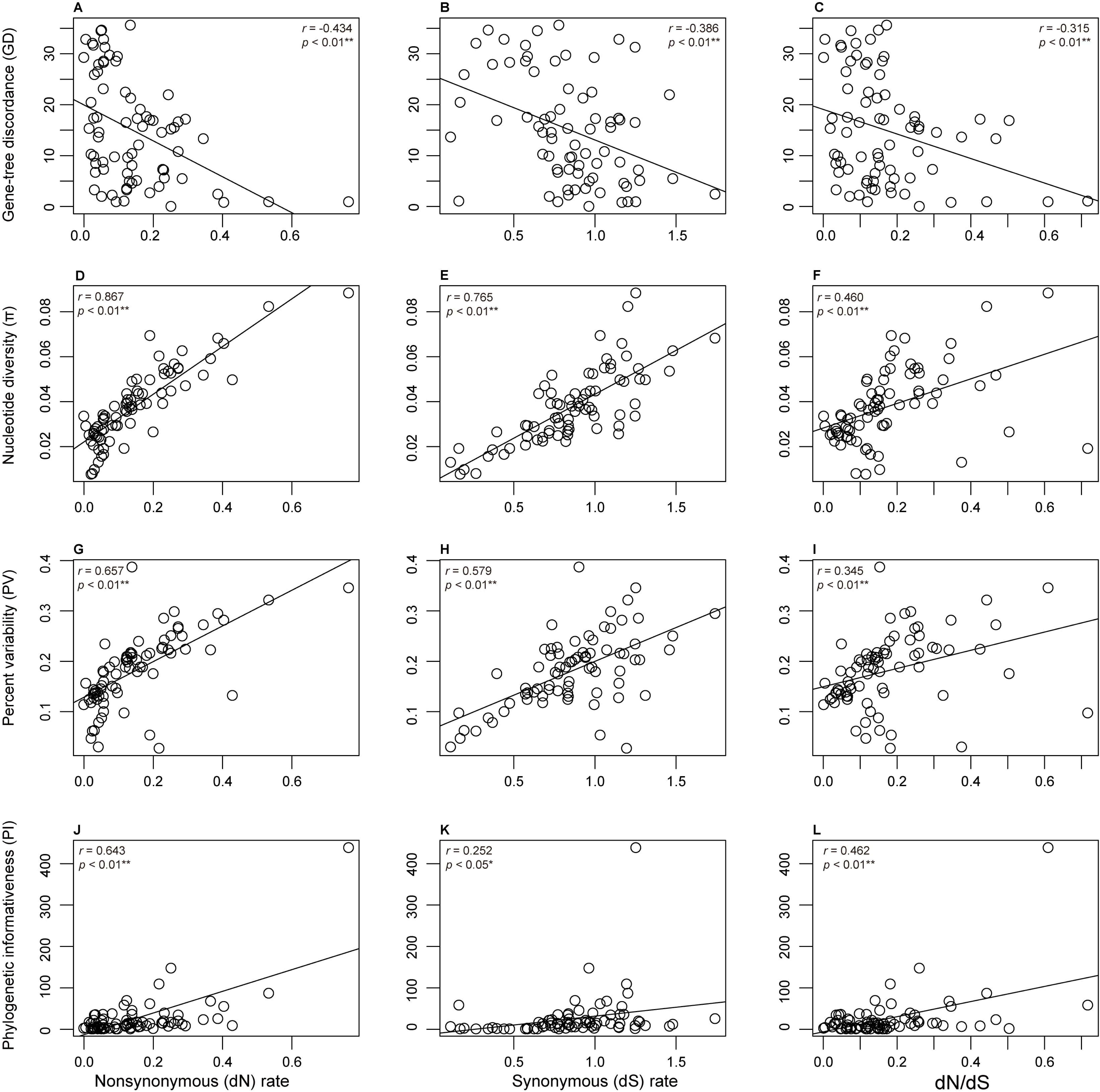
Correlation of evolutionary rate heterogeneity, including nonsynonymous (d*N*), synonymous (d*S*) substitution rates and d*N*/d*S*, with **(A-C)** gene tree discordance, **(D-F)** Nucleotide diversity (π), **(G-I)** percent variability (PV) and **(J-L)** phylogenetic informativeness. The stars next to the p-value were used to assess the level of significance.

### Correlation analysis

Correlation analysis showed all tested correlations were significant with a 0.05 cutoff. Specifically, nucleotide diversity (π), percent variability (PV) and phylogenetic informativeness (PI) were all positively correlated with the rates of nucleotide substitution, whereas gene-tree discordance (GD) was negatively correlated with the rates of nucleotide substitution.

## Discussion

### Phylogenetic implications of plastome-scale dataset

To our knowledge, the results presented here are the first to utilize a phylogenomic data set to investigate phylogenetic affinities among genera of Gentianeae, especially for the subtribe Swertiinae. Our study presents substantial improvements in tree resolution compared to previous phylogenetic reconstructions [36-40, 44], and provide a robust backbone of Gentianeae. Our phylogenomic backbone shows a clear subdivision of the Gentianeae into subtribes Gentianinae and Swertiinae, which is consistent with previous morphological [45] and molecular phylogenies [36-41, 44]. Gentianinae is consistently recognized as encompassing four genera --*Gentiana* L., *Tripterospermum* Blume, *Metagentiana* T.N.Ho & S.W.Liu, *Crawfurdia* Wall. Favre et al. [41] excluded *Gentiana* sect. *Otophora* from *Gentiana* and elevated as *Kuepferia* Adr.Favre and described *Sinogentiana* Adr.Favre & Y.M.Yuan by excluding two species from *Metagentiana*. Our results resolve *Gentiana* as monophyletic, and show a close relationship between *Kuepferia* and *Tripterospermum*, supporting the elevation of *Kuepferia*. Compared to Gentianinae, Swertiinae is more complicated due to the paraphyly of *Swertia* [38]. Our phylogenomic framework is congruent with the phylogeny of Swertiinae inferred using *trnL*-intron + *matK* [36, 37, 44], *atpB-rbcL* spacer [38], the supermatrix of eight plastid markers (*rbcL+ matK+ atpB+ ndhF+ rpl16+ rps16+ trnL-trnF* +*atpB-rbcL*) [40] and ITS [36-38, 44]. Overall, the present study places Gentianeae into a phylogenomic framework constituting the first steps in deeply understanding its evolutionary history. Further studies focusing on biogeography and diversification with denser sampling and more advanced methods are needed.

The majority of phylogenetic relationships of major groups of angiosperms that have been investigated in the last few decades rely mostly on ML inference of concatenated alignments of plastid genes (e.g. [7, 9, 10]). However, phylogenetic discordance produced by individual plastid genes has generally been largely ignored due to the fundamental assumption of a tightly linked unit of the plastome in coalescent theory. Goncalves et al. [18] showed that concatenated matrices may produce highly supported phylogenies that are discordant with individual gene trees. Walker et al. [14] demonstrated rampant gene-tree conflict within the plastome at all levels of angiosperm phylogeny, highlighting the necessity of future research into the consideration of plastome conflict. Both studies emphasized the importance of considering variation in phylogenetic signal across plastid genes and advocated the use of MSC with plastome matrices in phylogenomic investigations. In our analyses, despite the consistency between the tree topology produced by a concatenated matrix with that using MSC methods, gene tree topologies showed great variation with the species trees inferred from the concatenated data. Moreover, our computation of PCS revealed 23 of 76 plastid genes with negative scores and 20 genes with ambiguous estimation, indicating more than half of the genes are contributing to phylogenetic ambiguity of Gentianeae. A possible explanation for consistent topologies produced by the two methods is that the individual gene genealogy effect was too small to blur the accuracy of phylogenetic inference when all the genes were concatenated into a “supermatrix”. Methodologically, the individual gene trees and their bootstrap replicates that were used as inputs of MSC method in ASTRAL-III were inferred using ML in RAxML [46]. In such cases, conducting phylogenetic inference with concatenated genes as a single locus would represent a special case of MSC [25]. Nonetheless, such kind of gene-tree heterogeneity shouldn’t be disregarded, as it may influence divergence time estimation or higher taxonomic level phylogenetic reconstruction.

Our estimation of PCS scores revealed one (*atpB*) of the three widely used plastid markers (*rbcL, atpB* and *matK*) in phylogenetic inference of Gentianeae was an outlier gene that may have a disproportionate influence on the resolution of contentious relationships [43]. However, phylogenetic analysis of Gentianeae using a plastid supermatrix including the *atpB* gene [40] obtained a generally congruent tree topology with topologies from other markers [36-38]. A possible explanation for the observed consistency is that a concatenated matrix of *atpB* with other plastid genes may counteract the potential bias of *atpB* in the reconstruction of Gentianeae relationships. In addition, the relatively low PCS score of *matK* (PCS = 4) is a likely reason for extensive parallel clades existing in the study by Xi et al. (2014) [44].

### Gene characteristic diversity in plastomes of Gentianeae

Plastomes of Gentianeae are highly conserved in terms of genome structure with only one trivial IR expansion detected (Figure 1 and 2). Gene loss events involving *ndh* genes and *rpl33* in some species were detected. Loss of *ndh* genes in plastomes of *Gentiana* were previously reported in studies of Fu et al. [47] and Sun et al. [48]. In green plants, *ndh* genes encode components of the thylakoid ndh-complex involved in photosynthetic electron transport [49]. Recently, a comprehensive survey of gene loss and evolution of the plastomes showed *ndh* genes were the most commonly lost genes, suggesting that not all *ndh* genes are involved in or essential for functional electron transport [50]. Notably, the loss of *ndh* genes occurred within the genus *Gentiana* and formed a distinct clade, suggesting this loss may be related to adaptation of specific *Gentiana* species. Given that few plastomes of *Gentiana* are available compared to the total number of species (c. 360–400 species), further exploration with plastome sequencing is still required. In contrast to *ndh* genes, the loss of *rpl33* is likely more random. There is a stop codon in coding region of *rpl33* in *Comastoma pulmonarium* due to the change of cytosine (C) to thymine (T) at base pair 22. We also mapped all the sequenced reads of *C. pulmonarium* to the assembled sequence using Geneious for validation. Mapped reads showed almost all the reads supported the mutation. In the plastome of *Swertia hispidicalyx*, there is a small deletion containing the coding region of *rpl33* between the *psaJ* and *rps18* gene. A previous study suggested the loss of *rpl22, rpl32*, and *rpl33* genes was more prominent than the loss of other *rpl* genes [50]. In addition, a reverse genetics study found that knockout of the gene encoding ribosomal protein rpl33 did not affect plant viability and growth under standard conditions [51]. Hence, rpl33 may be a nonessential plastid ribosomal protein in plant photosynthesis, and loss of *rpl33* genes may be compensated for by other *rpl* genes or by nuclear encoded genes. Although the two gene loss events may not be related, they do reflect taxonomic diversity of the phylogeny by occurring in two distinct genera.

Estimating nucleotide substitution rates among different genes and different functional groups provided insight into the diverse selection regimes acting on plastomes evolution (e.g. [16, 20, 22]). Genes encoding subunits involved in photosynthetic processes, e.g. ATP synthase (ATP), NAD(P)H dehydrogenase (NDH), cytochrome b6f complex (PET), and photosystems I and II (PSA and PSB), have been shown to have lower rates of nucleotide substitution than other functional groups of genes in Poaceae [34] and Geraniaceae [17]. Similar patterns were observed in Gentianeae (Figure 4B). Among the photosynthetic functional groups, NDH had relatively higher nucleotide substitution rates, which was likely associated with the gene loss events in *Gentiana*. We identified a few functional gene groups that have accelerated substitution rates in Gentianeae, mainly the ribosomal protein (RPL and RPS) genes and the RNA polymerase (RPO) genes. Similar patterns have been previously documented, such as RPL and RPS genes were shown to be highly accelerated in Geraniaceae [20, 28] and RPO genes in Annonaceae, Passifloraceae and Geraniaceae [52]. In addition, *accD, clpP, ycf1*, and *ycf2* had the most accelerated rates [22] as detected in plastomes of *Silene* (Caryophyllaceae), and *clpP* and *ycf1* were found to have the highest d*N* values among genes in legumes [16]. In the present study, *ycf1* and *matK* had the highest d*N* value, and *ycf2* had the highest d*N*/d*S* value. Despite divergence of nucleotide substitution rates among individual genes or among functional groups of genes, there was no sign of positive selection acting on plastid genes of Gentianeae plastomes. Additionally, given that no obvious structure variation was detected, the pattern of substitution rate variation in Gentianeae may be attributed to the heterogeneity of genome-wide mutation rate. Furthermore, varied substitution rates in plastomes along with gene-tree discordance support the view that plastid genes are not tightly linked as previously thought and are experiencing different evolutionary forces [18].

Evolutionary dynamics of the plastid IR region has been previously documented [19]. A recent study demonstrated that synonymous substitution rates were, on average, 3.7 times slower in IR genes than in SC genes using 69 plastomes across 52 families of angiosperms, gymnosperms, and ferns [27]. However, study of *Pelargonium* (Geraniaceae) observed the opposite pattern in which d*S* values were higher for genes in the IR versus the SC regions [29]. Our results reveal d*S* rates about four to five times higher in LSC and SSC regions than IR region using a concatenated data set of genes in each region. The observed high d*S* value in the SSC region likely results from the six NDH genes involved in gene loss. In turn, high synonymous substitution rates may indicate relaxed selective constrains are responsible for the gene loss events. The low substitution rates in IR region is can be explained by the two identical copies providing a template for error correction when a mutation occurs in one of the copies [29].

### Correlation of evolutionary rate heterogeneity with gene characteristics and GD

Characteristics of plastid genes are of great importance in our understanding of plastome evolution and phylogenetic inference. We demonstrated extensive difference among plastid genes and functional groups of genes. The percent variability (PV) of PCGs exhibited similar pattern with nucleotide substitution rates of photosynthetic functional groups (ATP, NDH, PET, PSA and PSB) having lower values than ribosomal protein (RPL and RPS) and the RNA polymerase (RPO) groups, whereas the NDH group had higher values of nucleotide diversity (π). The value of π was estimated by the average number of nucleotide differences per site between two sequences and its sampling variance [53], and hence its estimation may be affected by the loss of *ndh* genes in *Gentiana*. Our results revealed a significant positive correlation of genetic variation with nucleotide substitution rates, indicating that diverse selection pressure is playing important roles in plastome evolution.

The net phylogenetic informativeness (PI) of a plastid gene reflects its performance in resolving complex phylogenetic relationships. Just one (*matK*) of the three (*matK, ndhB* and *rbcL*) markers widely used in phylogenetic studies of Gentianeae showed high net PI value, explaining the limited resolution in previous analyses and highlighting the utility of plastome-scale data sets. Among the genes tested, *ycf1* and *rpoC2* exhibited high net PI values, and accompanied by their relatively long gene length, would be optimal markers for phylogenetic inference of Gentianeae in the future. Indeed, the phylogenetic utility of *ycf1* has been demonstrated previously in orchids [54] as well as in radiating lineage [55], along with serving as a core barcode of land plants [56]. Our analyses revealed good performance of *rpoC2*, with high PI, lowest gene-tree discordance and positive high PCS score. Thus, we advocate *rpoC2* as a promising plastid DNA barcode for taxonomic study of Gentianeae, similar to the usefulness of *rpoC2* in the phylogenetic reconstruction of the angiosperm phylogeny [14].

We found a significant positive correlation of PI with nucleotide substitution rates, suggesting nucleotide substitutions of plastid genes are only slightly saturated. A sequence is considered saturated when it has undergone multiple substitutions that decreases phylogenetic information contained in the sequence due to underestimation of real genetic distance using the apparent distance [57]. Our analysis revealed negative correlation between gene-tree discordance (GD) and nucleotide substitution rates. Previous studies have drawn attention to the correlation between nucleotide substitution rates with number of indels and genomic rearrangements, such as in Geraniaceae [30], legumes [16, 21] and Lentibulariaceae [58], while GD remained poorly examined. As synonymous substitutions are neutral, changes in d*S* are likely to reflect changes in the underlying mutation rate, possibly due to problems with DNA repair, while non-synonymous (d*N*) substitutions rates and d*N*/d*S* are impacted not only by the underlying mutation rate, but also by selection. Given no sign of positive selection among plastid genes of Gentianeae, the correlation between GD with d*N* and d*N*/d*S* suggested GD is likely governed by the strength of purifying selection or the selective removal of deleterious mutations. The negative correlation between GD and d*N*/d*S* indicates that gene-tree discordance is more rampant under higher strength of purifying selection. Selection against deleterious mutations may cause a reduction in the amount of genetic variability at linked neutral sites [59], and hence rapid removal of mutations may blur the evolutionary footprints of a lineage.

## Conclusions

Our results presented here are the first to utilize a phylogenomic data set to investigate phylogenetic relationships among genera of Gentianeae. The phylogenomic framework lays the foundation for deep understanding of the evolutionary history of this diverse tribe. Comparative genomic analyses reveal both extensive evolutionary rate heterogeneity and genetic variation among plastid genes, supporting the view that plastid genes are not tightly linked as previously thought and are experiencing different evolutionary forces. Of the commonly used markers in phylogenetic inference of Gentianeae, only *matK* has high phylogenetic informativeness, while *atpB* may have a disproportionate influence on the resolution of contentious relationships. The rarely used gene *rpoC2* is the top-performing gene, similar to the usefulness in the phylogenetic reconstruction of the angiosperm phylogeny, and hence is advocated as a promising plastid DNA barcode for taxonomic studies of Gentianeae. Notably, the rampant phylogenetic discordance of gene tree was detected, highlighting the necessity of considering gene-tree heterogeneity into future phylogenomic studies.

## Methods

### Taxon sampling and sequencing

Fresh leaves of ten species of Gentianeae were collected from the Qinghai-Tibet Plateau (QTP) and adjacent regions complying with national guidelines. Voucher specimens were identified by the Herbarium of Kunming Institute of Botany (KUN) and deposited there. Sample information is provided in supplementary materials Table S1. For all species, total genomic DNA was extracted following the procedure of Plant Genomic DNA Kit (DP305) from Tiangen Biotech (Beijing) Co., Ltd., China. Illumina paired-end libraries were constructed with the NEBNext® Ultra™ DNA Library Prep Kit (New England Biolabs, Ipswich, MA, USA) according to the manufacturer’s protocol. A 500-bp DNA TruSeq Illumina (Illumina Inc., San Diego, CA, USA) sequencing library was constructed using 2.5-5.0 ng sonicated DNA as input and final quantifications were done using an Agilent 2100 Bioanalyzer (Agilent Technologies, Santa Clara, CA, USA) and real-time quantitative PCR. Libraries were multiplexed and sequenced using a 2×150 bp run on an Illumina HiSeq 4000 platform at Novogene Co., Ltd in Kunming, Yunnan, China.

### Plastome assembly and annotation

Raw sequence reads were filtered using Trimmomatic v.0.36 [60] by removing duplicate reads, adapter-contaminated reads, and reads with more than five Ns. Remaining high-quality reads were assembled *de novo* into contigs in NOVOPlasty v.2.7.2 [61] using a seed-and-extend algorithm with the plastome sequence of *Swertia mussotii* (Genbank accession: NC_031155.1) as the seed input and other parameters remaining at default settings (see NOVOPlasty manual). Assembled plastomes were annotated using Plastid Genome Annotator (PGA) [62]. Start/stop codons and intron/exon boundaries were determined based on published plastomes of Gentianeae as a reference in Geneious v.9.0.5 [63]. The tRNA genes were identified with tRNAscan-SE [64]. For comparation, a linear graphical map of all sequenced plastomes were visualized with OGDRAW [65].

### Phylogenetic analyses

Twenty-nine taxa of Gentianeae (17 Gentianinae + 12 Swertiinae) and three outgroups (*Coffea arabica, Coffea canephora* and *Galium aparine*) were included in phylogenomic analyses (Table S1). Both concatenated and coalescent analyses were conducted. For the concatenated approach, the common 76 protein coding genes (PCGs) were extracted using PhyloSuite v.1.1.15 [66] with subsequent manual modifications. The sequences of the 76 PCGs were aligned in batches with MAFFT v.7.313 using “G-INS-i” strategy and codon alignment mode, and then concatenated in PhyloSuite. Both partitioned and unpartitioned analyses were performed. For data partitioning, PartitionFinder v.1.0.1 [67] was implemented to determine optimal partitioning scheme and evolutionary model selection under the Bayesian Information Criterion (BIC). Maximum likelihood (ML) analysis was conducted in RAxML v.8.2.10 [68] under the “GTRGAMMA” model with the “rapid bootstrap” algorithm (1000 replicates). Bayesian inference (BI) was performed using MrBayes v.3.2.3 [69] with four Markov chains (one cold and three heated) running for 5,000,000 generations from a random starting tree and sampled every 5000 generations. The first 25% of the trees were discarded as burn-in, and the remaining trees were used to construct majority-rule consensus trees.

For the coalescent analysis, ML unrooted trees for each individual gene were estimated separately using RAxML under the “GTRGAMMA” model with 500 bootstrap replicates. ASTRAL-III v.5.6.2 [46] algorithm was used to estimate the species tree from 76 gene trees with node supports calculated as local posterior probabilities.

### Exploration of plastid gene tree landscape

To explore gene tree variations, we plotted the statistical distribution of trees with Robinson-Foulds algorithms [70], by calculating distances between unrooted trees using the R package TREESPACE v.1.0.0 [42] following the workflow of Goncalves et al. [18], and visualizing with ggplot2 v.2.2.1 [71]. Since TREESPACE only accepts groups of trees containing the same tips, we removed the *rpl33* locus which was absent in some species. Additionally, we removed six species of *Gentiana* from analysis due to the loss of *ndh* genes: *G. hexaphylla, G. oreodoxa, G. veitchiorum, G. ornate* and *G. caelestis*. We also included two species trees inferred from concatenated and coalescent analyses. In total, the dataset consisted of 75 gene trees from 26 taxa and two species trees. We used the distance between gene trees and the coalescent species tree to estimate gene-tree discordance (GD) of plastid genes. Distances were calculated using the first two PCoAs estimated by TREESPACE.

In addition, partitioned coalescence support (PCS) was calculated for each PCG using scripts provided by Gatesy et al. [43] (https://github.com/dbsloan). PCS quantifies the positive or negative influence of each gene tree in a phylogenomic data set for clades supported by summary coalescence methods [43, 72]. We used the phylogenetic tree generated by ASTRAL-III as the optimal species tree topology, and the tree inferred by RAxML as an alternative species tree.

### Nucleotide substitution rate analysis

To estimate rates of nucleotide substitution, nonsynonymous (d*N*), synonymous (d*S*), and the ratio of nonsynonymous to synonymous rates (d*N*/d*S*) were calculated in PAML v.4.9 [73] using the CODEML option with codon frequencies estimated using the F3 × 4 model. The phylogeny generated using the concatenated method was used as a constraint tree. Gapped regions were removed using “cleandata = 1” option. The “model=0” option was used for allowing a single d*N*/d*S* value to vary among branches. Other parameters in the CODEML control file were left at default settings. For the comparisons between different functional groups of PCGs, we consolidated the 76 PCGs into nine groups, i.e. photosystem I (PSA), photosystem II (PSB), cytochrome B6f complex (PET), ATP synthase (ATP), rubisco large subunit (Rubisco), RNA polymerase (RPO), ribosomal proteins large subunit (RPL), ribosomal proteins small subunit (RPS) and NADH dehydrogenase (NDH), and other genes (OG, including *ycf1, ycf2, ycf3, ycf4, accD, clpP, ccsA, cemA* and *matK*). Detailed information of functional groups is provided in Table 2.

### Genetic variation and phylogenetic informativeness of PCGs

We used nucleotide diversity (π) and the percent variability (PV) to represent genetic variation of PCGs. Percent variability of each PCG was estimated by dividing segregating sites (S, i.e. the number of variable positions) by the length of gene. Both π and S were calculated in the program DnaSP v.6.0.7 [74] using sequences of the 76 PCGs aligned by MAFFT separately. The PhyDesign web application (http://phydesign.townsend.yale.edu/) [75] was used to estimate the net phylogenetic informativeness (PI) profiles for the 76 PCGs by using the HyPhy substitution rates algorithm for DNA sequences [76]. A relative-time ultrametric tree constructed in the *dnamlk* program in PHYLIP [77] based on ML tree, along with the 76 PCGs alignment partitioned by genes were used as input files in PhyDesign to calculate phylogenetic informativeness.

### Correlation analysis

Correlation of nucleotide substitution rates (including d*N*, d*S* and d*N*/d*S* values of each gene) with gene-tree discordance (GD), nucleotide diversity (π), percent variability (PV) and phylogenetic informativeness (PI) were tested using *cor* function in R package *stats* v3.6.1 (https://www.rdocumentation.org/packages/stats). The function *cor*.*test* was used for calculating values with Pearson test.

## Authors’ contributions

HCW and HS conceived and designed the study. XZ and YXS performed de novo assembly, genome annotation, phylogenetic and other analyses. XZ, YXS, JBL, HCW and HS drafted the manuscript. JS and HJZ performed the experiments. XZ, ZYL, LLJ, JS, NL and TD collected the leaf materials. All authors discussed the results and helped shape the research, analyses and final manuscript.

## Figure legends

**Figure S1** Loss of *rpl33* coding region in plastomes of *Comastoma pulmonarium* and *Swertia hispidicalyx*. There is a stop codon in coding region of *rpl33* in *Comastoma pulmonarium* due to the change of cytosine (C) to thymine (T) at 22bp, and a small deletion containing the coding region of *rpl33* between the *psaJ* and *rps18* gene in the plastome of *Swertia hispidicalyx*.

**Figure S2** Maximum likelihood phylogram of Gentianeae from unpartitioned concatenated matrix of 76 plastid protein-encoding genes using RAxML. Maximum likelihood bootstrap (BS) values and the PP calculated from MrBayes are shown at nodes, except nodes with 100% BS and 1.0 PP.

**Figure S3** Phylogenetic informativeness profile estimated in PhyDesign. **(A)** The ultramteric tree of Gentianeae. **(B)** Net phylogenetic informativeness profile for 76 plastid protein-coding genes. Ten genes with the greatest informativeness are color-coded and indicated at the right. X-and Y-axes represent relative-time and net phylogenetic informativeness, respectively.

