## Supplemental Figures S1-S3 for "Plastome phylogenomic study of Gentianeae (Gentianaceae): widespread gene tree discordance and its association with evolutionary rate heterogeneity of plastid genes"

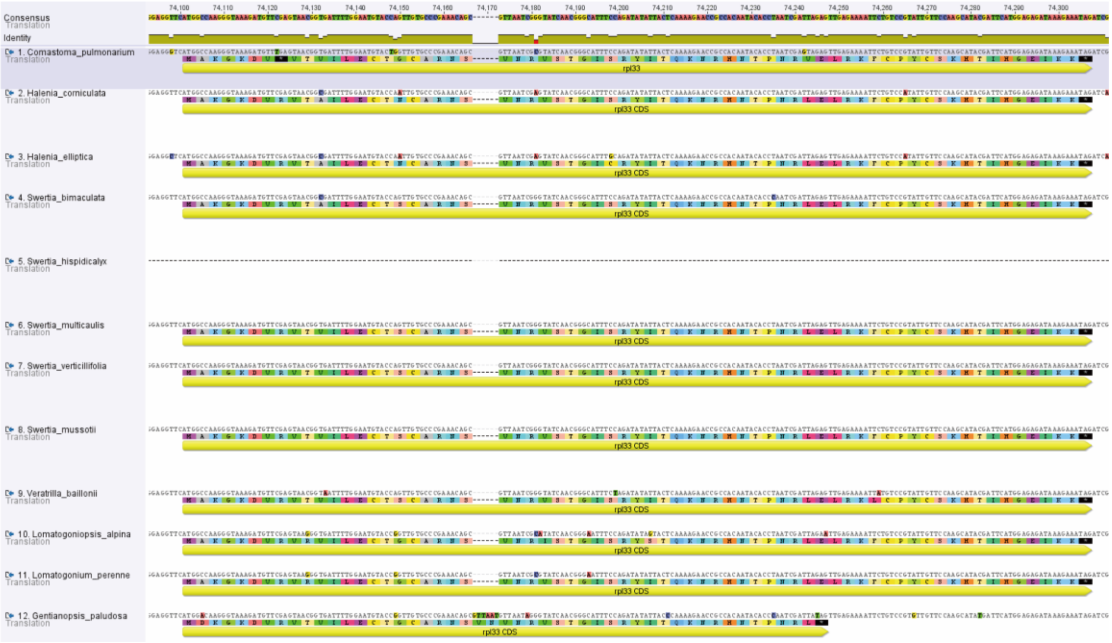


**Figure S1** Loss of *rpl33* coding region in plastomes of *Comastoma pulmonarium and Swertia hispidicalyx*. There is a stop codon in coding region of *rpl33* in *Comastoma pulmonarium* due to the change of cytosine (C) to thymine (T) at 22bp, and a small deletion containing the coding region of rpl33 between the *psaJ* and *rps18* gene in the plastome of *Swertia hispidicalyx*.

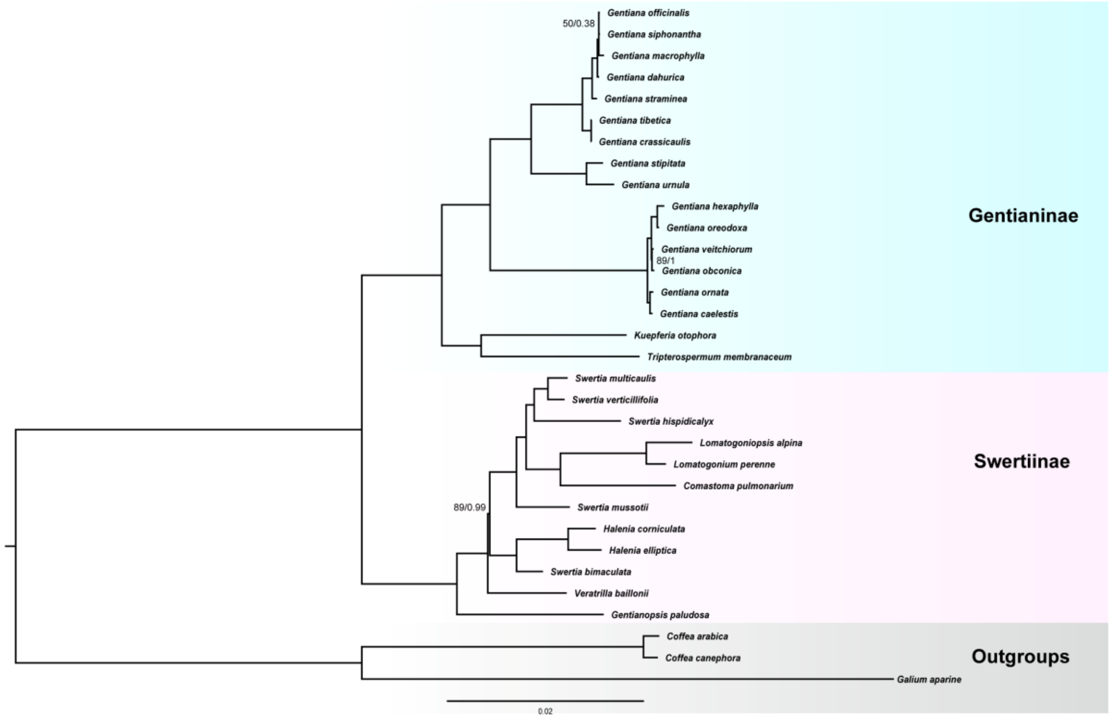


**Figure S2** Maximum likelihood phylogram of Gentianeae from unpartitioned concatenated matrix of 76 plastid protein-encoding genes using RAxML. Maximum likelihood bootstrap (BS) values and the PP calculated from MrBayes are shown at nodes, except nodes with 100% BS and 1.0 PP.

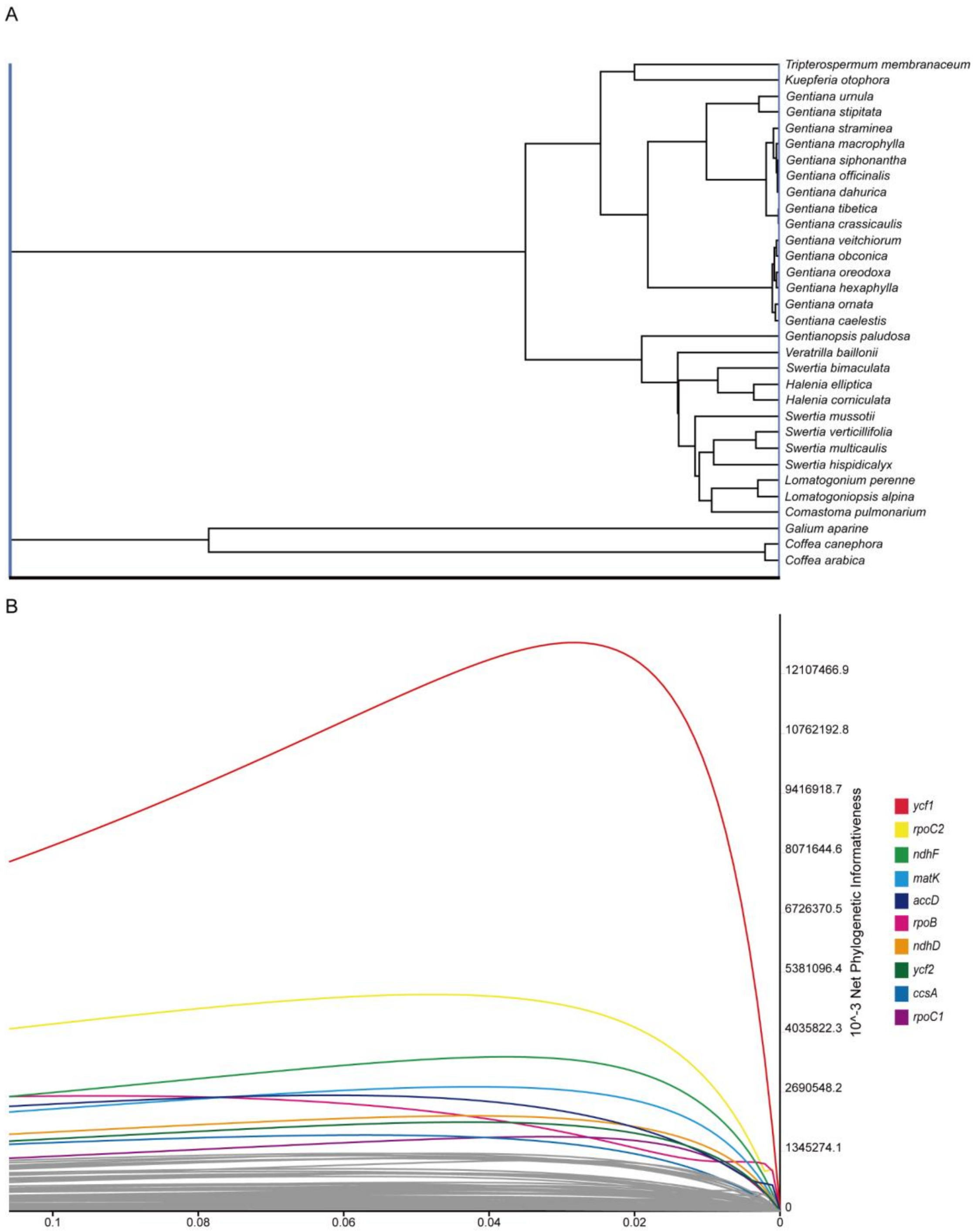


**Figure S3** Phylogenetic informativeness profile estimated in PhyDesign. (A) The ultramteric tree of Gentianeae. (B) Net phylogenetic informativeness profile for 76 plastid protein-coding genes. Ten genes with the greatest informativeness are color-coded and indicated at the right. X- and Y-axes represent relative-time and net phylogenetic informativeness, respectively.
