## Supplementary material for "Plastome phylogenomic study of Gentianeae (Gentianaceae): widespread gene tree discordance and its association with evolutionary rate heterogeneity of plastid genes": Table S1-S2

**Table S1** Taxa included in present study. NCBI accession numbers and voucher specimens’ information are provided for newly sequenced plastomes, “-” indicates no applicable

| **Family** | **Subtribes** | **Species** | **NCBI accession numbers** | **Voucher specimens information** |
| --- | --- | --- | --- | --- |
| Gentianaceae | Swertiinae | *Comastoma pulmonarium* | MT228723 | FSC-342 |
| Gentianaceae | Swertiinae | *Gentiana urnula* | MT228724 | ZJW6386 |
| Gentianaceae | Swertiinae | *Gentianopsis paludosa* | MT228725 | FSC-55 |
| Gentianaceae | Swertiinae | *Halenia elliptica* | MT228726 | FSC-339 |
| Gentianaceae | Swertiinae | *Lomatogoniopsis alpina* | MT228728 | ZJW5972 |
| Gentianaceae | Swertiinae | *Lomatogonium perenne* | MT228729 | ZJW5118 |
| Gentianaceae | Swertiinae | *Swertia multicaulis* | MT228730 | ZJW5106 |
| Gentianaceae | Swertiinae | *Veratrilla baillonii* | MT228732 | ZJW6591 |
| Gentianaceae | Swertiinae | *Halenia corniculata* | MK606372.1 | - |
| Gentianaceae | Swertiinae | *Swertia hispidicalyx* | MH321887.1 | - |
| Gentianaceae | Swertiinae | *Swertia mussotii* | KU641021.1 | - |
| Gentianaceae | Swertiinae | *Swertia verticillifolia* | MF795137 | - |
| Gentianaceae | Swertiinae | *Swertia bimaculata* | MH394374 | - |
| Gentianaceae | Gentianinae | *Kuepferia otophora* | MT228727 | FSC-76 |
| Gentianaceae | Gentianinae | *Tripterospermum membranaceum* | MT228731 | KUN1220748 |
| Gentianaceae | Gentianinae | *Gentiana caelestis* | MG192304.1 | - |
| Gentianaceae | Gentianinae | *Gentiana crassicaulis* | KJ676538.1 | - |
| Gentianaceae | Gentianinae | *Gentiana dahurica* | MH261259.1 | - |
| Gentianaceae | Gentianinae | *Gentiana hexaphylla* | MG192305.1 | - |
| Gentianaceae | Gentianinae | *Gentiana macrophylla* | KY856959.1 | - |
| Gentianaceae | Gentianinae | *Gentiana obconica* | MG192306.1 | - |
| Gentianaceae | Gentianinae | *Gentiana officinalis* | MH261261.1 | - |
| Gentianaceae | Gentianinae | *Gentiana oreodoxa* | MG192307.1 | - |
| Gentianaceae | Gentianinae | *Gentiana ornata* | MG192308.1 | - |
| Gentianaceae | Gentianinae | *Gentiana siphonantha* | MH261260.1 | - |
| Gentianaceae | Gentianinae | *Gentiana stipitata* | MG192309.1 | - |
| Gentianaceae | Gentianinae | *Gentiana straminea* | KJ657732.1 | - |
| Gentianaceae | Gentianinae | *Gentiana tibetica* | KU975374.1 | - |
| Gentianaceae | Gentianinae | *Gentiana veitchiorum* | MG192310.1 | - |
| Rubiaceae | - | *Coffea arabica* | NC_008535.1 | - |
| Rubiaceae | - | *Coffea canephora* | KU500324.1 | - |
| Rubiaceae | - | *Galium aparine* | KY562587.1 | - |

**Table S2** Genetic characteristics of 76 protein-coding genes used in analyses, including nonsynonymous rate (d*N*), Synonymous rate (d*S*), nucleotide diversity (π), percent variability (PV), phylogenetic informativeness (PI), gene-tree discordance (GD) and partitioned coalescence support (PCS). Detailed information of functional group is provided in Table 2

| **Genes** | **Functional groups** | **d*N*** | **d*S*** | **d*N*/d*S*** | **π** | **PV** | **PI** | **GD** | **PCS** |
| --- | --- | --- | --- | --- | --- | --- | --- | --- | --- |
| *accD* | OG | 0.3653 | 1.0710 | 0.3411 | 0.0590 | 0.2228 | 67.5645 | 2.1426 | 52 |
| *atpA* | ATP | 0.0551 | 1.1528 | 0.0478 | 0.0343 | 0.1808 | 36.1631 | 8.7252 | -56 |
| *atpB* | ATP | 0.0514 | 0.7176 | 0.0716 | 0.0270 | 0.1514 | 21.5446 | 2.0286 | -56 |
| *atpE* | ATP | 0.1797 | 0.7273 | 0.2471 | 0.0392 | 0.2114 | 8.7099 | 17.6664 | 2 |
| *atpF* | ATP | 0.2248 | 0.7315 | 0.3073 | 0.0438 | 0.2250 | 15.2736 | 14.5917 | 27 |
| *atpH* | ATP | 0.0056 | 1.1477 | 0.0049 | 0.0291 | 0.1564 | 4.5894 | 32.8122 | 52 |
| *atpI* | ATP | 0.0924 | 0.7713 | 0.1197 | 0.0331 | 0.1748 | 13.1140 | 0.9808 | 48 |
| *ccsA* | OG | 0.4041 | 1.1645 | 0.3470 | 0.0658 | 0.2811 | 55.2155 | 0.8087 | 56 |
| *cemA* | OG | 0.3444 | 0.7356 | 0.4682 | 0.0517 | 0.2727 | 23.1032 | 13.3351 | 27 |
| *clpP* | OG | 0.1693 | 0.6523 | 0.2596 | 0.0436 | 0.1881 | 17.6351 | 15.7227 | 0 |
| *matK* | OG | 0.5333 | 1.2032 | 0.4433 | 0.0824 | 0.3216 | 87.6416 | 0.9279 | 4 |
| *ndhA* | NDH | 0.232 | 0.9876 | 0.2349 | 0.0524 | 0.2423 | 38.8263 | 5.5621 | 0 |
| *ndhB* | NDH | 0.0412 | 0.1099 | 0.3748 | 0.0129 | 0.0301 | 6.7568 | 13.6284 | 0 |
| *ndhC* | NDH | 0.1199 | 0.9808 | 0.1223 | 0.0361 | 0.1889 | 7.9042 | 22.433 | 0 |
| *ndhD* | NDH | 0.1904 | 1.0327 | 0.1844 | 0.0694 | 0.0537 | 62.1701 | 2.6858 | 0 |
| *ndhE* | NDH | 0.1535 | 1.0038 | 0.1530 | 0.0448 | 0.1799 | 9.7540 | 17.3181 | 0 |
| *ndhF* | NDH | 0.2168 | 1.1928 | 0.1818 | 0.0603 | 0.0271 | 109.9486 | 3.9719 | 0 |
| *ndhG* | NDH | 0.2711 | 1.0581 | 0.2562 | 0.0551 | 0.2689 | 19.9161 | 10.7515 | 0 |
| *ndhH* | NDH | 0.1388 | 1.1791 | 0.1177 | 0.0490 | 0.2152 | 36.1738 | 4.5341 | 0 |
| *ndhI* | NDH | 0.1511 | 1.2747 | 0.1185 | 0.0501 | 0.2032 | 17.8783 | 5.1046 | 0 |
| *ndhJ* | NDH | 0.1261 | 0.8440 | 0.1494 | 0.0410 | 0.1983 | 11.5506 | 9.5790 | 0 |
| *ndhK* | NDH | 0.1310 | 0.9647 | 0.1358 | 0.0376 | 0.1856 | 15.4820 | 4.9396 | 0 |
| *petA* | PET | 0.1236 | 0.9438 | 0.1309 | 0.0436 | 0.2090 | 21.9469 | 3.5395 | -56 |
| *petB* | PET | 0.0344 | 1.0109 | 0.0340 | 0.0279 | 0.1376 | 14.6061 | 8.4695 | 2 |
| *petD* | PET | 0.0554 | 0.8376 | 0.0662 | 0.0321 | 0.1303 | 10.2926 | 23.1140 | -52 |
| *petG* | PET | 0.0270 | 0.5722 | 0.0473 | 0.0207 | 0.1351 | 1.1984 | 31.7335 | 8 |
| *petL* | PET | 0.1336 | 0.7772 | 0.1719 | 0.0303 | 0.2151 | 1.3856 | 35.6260 | 0 |
| *petN* | PET | 0.0961 | 0.5843 | 0.1645 | 0.0293 | 0.1379 | 1.5554 | 29.4858 | -24 |
| *psaA* | PSA | 0.0326 | 0.7722 | 0.0422 | 0.0249 | 0.1409 | 35.2175 | 6.7164 | 0 |
| *psaB* | PSA | 0.0292 | 0.8319 | 0.0351 | 0.0268 | 0.1367 | 34.2358 | 3.2928 | -56 |
| *psaC* | PSA | 0.0273 | 1.1449 | 0.0238 | 0.0257 | 0.1276 | 5.2226 | 17.3406 | 54 |
| *psaI* | PSA | 0.1990 | 0.3948 | 0.5040 | 0.0266 | 0.1759 | 1.8664 | 16.8629 | -27 |
| *psaJ* | PSA | 0.0566 | 0.7561 | 0.0748 | 0.0326 | 0.1778 | 4.3983 | 28.5362 | 30 |
| *psbA* | PSB | 0.0151 | 0.8314 | 0.0182 | 0.0252 | 0.1246 | 16.5847 | 15.3867 | 28 |
| *psbB* | PSB | 0.0569 | 0.8398 | 0.0678 | 0.0278 | 0.1594 | 25.213 | 7.2645 | 1 |
| *psbC* | PSB | 0.0281 | 0.7164 | 0.0393 | 0.0260 | 0.1438 | 21.5323 | 9.8471 | 28 |
| *psbD* | PSB | 0.0205 | 0.6772 | 0.0303 | 0.0223 | 0.118 | 14.3902 | 10.2727 | -56 |
| *psbE* | PSB | 0.0376 | 0.5819 | 0.0646 | 0.0244 | 0.1245 | 2.8053 | 17.5954 | -25 |
| *psbF* | PSB | 0.0484 | 0.6442 | 0.0751 | 0.0229 | 0.1453 | 1.5568 | 34.5233 | -2 |
| *psbH* | PSB | 0.2929 | 0.6900 | 0.4245 | 0.0471 | 0.2237 | 8.1256 | 17.1520 | 56 |
| *psbI* | PSB | 0.0388 | 0.6232 | 0.0622 | 0.0288 | 0.1389 | 1.4368 | 26.5306 | 54 |
| *psbJ* | PSB | 0.0565 | 0.4395 | 0.1286 | 0.0165 | 0.1000 | 1.2624 | 32.8013 | 53 |
| *psbL* | PSB | 0.0509 | 0.3419 | 0.1487 | 0.0156 | 0.0877 | 1.1146 | 34.6862 | -27 |
| *psbM* | PSB | 0.0917 | 0.5728 | 0.1601 | 0.0295 | 0.1471 | 1.3478 | 28.6185 | 26 |
| *psbN* | PSB | 0.0565 | 0.4743 | 0.1191 | 0.0192 | 0.1163 | 1.4453 | 28.3032 | 2 |
| *psbT* | PSB | 0.0001 | 0.9968 | 0.0001 | 0.0336 | 0.1143 | 2.4292 | 29.2809 | 48 |
| *psbZ* | PSB | 0.0737 | 0.8185 | 0.0900 | 0.0223 | 0.1882 | 2.4807 | 29.7000 | 0 |
| *rbcL* | Rubisco | 0.0816 | 0.8366 | 0.0976 | 0.0294 | 0.1505 | 37.4035 | 2.2135 | 56 |
| *rpl14* | RPL | 0.0878 | 0.8567 | 0.1025 | 0.0380 | 0.1995 | 5.4226 | 9.7891 | -52 |
| *rpl16* | RPL | 0.1220 | 1.2469 | 0.0978 | 0.0390 | 0.2034 | 9.1429 | 16.4541 | 55 |
| *rpl2* | RPL | 0.0295 | 0.1926 | 0.1529 | 0.0098 | 0.0624 | 3.0633 | 25.9169 | 50 |
| *rpl20* | RPL | 0.2510 | 0.9705 | 0.2586 | 0.0449 | 0.2164 | 16.7373 | 15.1967 | 0 |
| *rpl22* | RPL | 0.3859 | 1.7402 | 0.2218 | 0.0683 | 0.2952 | 25.7191 | 2.4413 | 4 |
| *rpl23* | RPL | 0.0236 | 0.2666 | 0.0885 | 0.0079 | 0.0609 | 0.9822 | 32.11 | -26 |
| *rpl32* | RPL | 0.2438 | 1.4596 | 0.1670 | 0.0535 | 0.2222 | 7.6006 | 21.9948 | 28 |
| *rpl33* | RPL | 0.4289 | 1.3109 | 0.3249 | 0.0498 | 0.1324 | 9.2624 | NA | -24 |
| *rpl36* | RPL | 0.0609 | 1.2482 | 0.0488 | 0.0335 | 0.2342 | 1.7339 | 31.2538 | 54 |
| *rpoA* | RPO | 0.2720 | 1.0983 | 0.2476 | 0.0547 | 0.2653 | 34.6309 | 16.6631 | -56 |
| *rpoB* | RPO | 0.1229 | 0.8775 | 0.1401 | 0.0357 | 0.2036 | 69.2034 | 3.2748 | -28 |
| *rpoC1* | RPO | 0.1382 | 0.9017 | 0.1533 | 0.0381 | 0.3872 | 46.1063 | 8.1058 | -56 |
| *rpoC2* | RPO | 0.2508 | 0.9633 | 0.2603 | 0.0526 | 0.2509 | 147.8433 | 0 | 56 |
| *rps11* | RPS | 0.1891 | 1.154 | 0.1638 | 0.0498 | 0.2160 | 13.9206 | 17.0221 | 56 |
| *rps12* | RPS | 0.0423 | 0.3696 | 0.1144 | 0.0187 | 0.0780 | 4.0176 | 27.9498 | -56 |
| *rps14* | RPS | 0.158 | 0.8777 | 0.1800 | 0.0435 | 0.2400 | 5.9692 | 12.1429 | 0 |
| *rps15* | RPS | 0.2841 | 1.4808 | 0.1919 | 0.0627 | 0.2500 | 11.7144 | 5.4910 | 0 |
| *rps18* | RPS | 0.1622 | 0.7838 | 0.2070 | 0.0386 | 0.1883 | 6.5995 | 19.0925 | -28 |
| *rps19* | RPS | 0.1347 | 0.9256 | 0.1456 | 0.0408 | 0.2174 | 4.8625 | 21.3507 | 56 |
| *rps2* | RPS | 0.1390 | 0.9439 | 0.1472 | 0.0365 | 0.2076 | 15.7626 | 10.3947 | 0 |
| *rps3* | RPS | 0.2297 | 1.2682 | 0.1811 | 0.0548 | 0.2861 | 19.7711 | 7.1380 | 27 |
| *rps4* | RPS | 0.1255 | 0.8958 | 0.1401 | 0.0394 | 0.2085 | 13.7031 | 6.4906 | 0 |
| *rps7* | RPS | 0.0196 | 0.1704 | 0.1152 | 0.0077 | 0.0473 | 1.2825 | 20.4554 | 50 |
| *rps8* | RPS | 0.2600 | 1.0960 | 0.2372 | 0.0568 | 0.2990 | 13.6557 | 15.5247 | -24 |
| *ycf1* | OG | 0.7634 | 1.2515 | 0.6100 | 0.0884 | 0.3460 | 438.8369 | 0.9808 | -56 |
| *ycf2* | OG | 0.1163 | 0.1621 | 0.7173 | 0.0191 | 0.0974 | 58.8030 | 1.0214 | -28 |
| *ycf3* | OG | 0.0429 | 0.6762 | 0.0635 | 0.0243 | 0.1310 | 6.1739 | 14.5601 | 4 |
| *ycf4* | OG | 0.2270 | 0.7662 | 0.2963 | 0.0393 | 0.2283 | 14.4875 | 7.2522 | -52 |
